# Breakthrough infection elicits hypermutated IGHV3-53/3-66 public antibodies with broad and potent neutralizing activity against SARS-CoV-2 variants including BQ and XBB lineages

**DOI:** 10.1101/2023.07.18.549524

**Authors:** Ling Li, Xixian Chen, Zuowei Wang, Yunjian Li, Chen Wang, Liwei Jiang, Teng Zuo

**Affiliations:** Laboratory of Immunoengineering, Institute of Health and Medical Technology, Hefei Institutes of Physical Science, Chinese Academy of Sciences, Hefei 230031, China; University of Science and Technology of China, Hefei 230026, China

## Abstract

The rapid emergence of SARS-CoV-2 variants of concern (VOCs) calls for efforts to study broadly neutralizing antibodies elicited by infection or vaccination so as to inform the development of vaccines and antibody therapeutics with broad protection. Here, we identified two convalescents of breakthrough infection with relatively high neutralizing titers against all tested viruses including BQ and XBB lineages. Among 50 spike-specific monoclonal antibodies (mAbs) cloned from their B cells, the top 6 neutralizing mAbs (KXD01-06) belong to previously defined IGHV3-53/3-66 public antibodies. Although most antibodies in this class are dramatically escaped by VOCs, KXD01-06 exhibit broad neutralizing capacity with the IC50s of KXD01 ranging from 0.011∼0.059μg/ml. Deep mutational scanning reveals that KXD01-06 target highly conserved sites on RBD including D420, Y421, L455, F456, A475 and N487. Genetic and functional analysis further indicates that the extent of somatic hypermutation is critical for the breadth of IGHV3-53/3-66 public antibodies. Overall, we discovered and characterized IGHV3-53/3-66 public antibodies with broad and potent neutralizing activity against SARS-CoV-2, which provides rationale for novel vaccines and antibody therapeutics based on this class of antibodies.

## Introduction

Since the outbreak of pandemic caused by SARS-CoV-2, numerous successes have been achieved in the development of vaccines and antibody therapeutics. Hundreds of vaccine candidates have been developed and 50 vaccines have been approved as of December 2022 (https://covid19.trackvaccines.org/). Meanwhile, tens of thousands of neutralizing monoclonal antibodies (mAbs) have been identified and 11 mAbs have successively received emergency use authorization within the first two years of the pandemic^1^. However, these successes have been gradually destroyed by the continuous emergence of SARS-CoV-2 variants. According to the transmissibility, virulence and immune escape of variants, the World Health Organization (WHO) has identified a succession of variants of concern (VOCs), including Alpha (B.1.1.7), Beta (B.1.351), Gamma (P.1), Delta (B.1.617.2) and Omicron (B.1.1.529). Omicron was first reported in South Africa in November 2021 and quickly became predominant worldwide^2^. To date, Omicron has developed hundreds of subvariants, which are classified into five main lineages including BA.1, BA.2, BA.3, BA.4, BA.5 and a serial of sublineages such as BA.1.1, BA.2.12.1, BA.2.75, BA.4.6, BF.7, BQ.1 and XBB^3^. Compared with SARS-CoV-2 prototype and earlier VOCs, Omicron lineages, particularly the emerging BQ and XBB sublineages, display most striking neutralization evasion from infection– and vaccination-elicited polyclonal antibodies and previously identified mAbs^4–8^. The persistent evolution of SARS-CoV-2 highlights the urgency to develop vaccines and antibody therapeutics with broad protection against current and future SARS-CoV-2 variants.

Spike (S) protein on virus surface mediates viral entry into host cells and is therefore the main target of vaccines and antibody therapeutics. Structurally, S protein is trimeric and each monomer consists of two functional subunits: S1 for engaging the receptor angiotensin converting enzyme 2 (ACE2) and S2 for driving fusion of viral and cellular membranes. S1 contains an amino-terminal (N-terminal) domain (NTD), a receptor-binding domain (RBD) and two carboxy-terminal (C-terminal) domains (CTD1 and CTD2). RBD further consists of a core structure and a receptor binding motif (RBM) that contacts with ACE2. S2 includes the N-terminal fusion peptide and its proximity region, heptad repeat 1 (HR1), central helix, stem helix, HR2, transmembrane region, and cytoplasmic tail. Although all the extracellular domains are susceptible to antibody binding, the majority of neutralizing antibodies target RBD with the rest recognizing NTD, S2, CTD or other epitopes^9, 10^.

A large-scale V gene usage analysis has revealed that RBD-specific antibodies are most frequently encoded by IGHV3-53 and the highly related IGHV3-66, and therefore these antibodies are termed as IGHV3-53/3-66 public antibodies^11^. Exemplified by CB6 (LY-CoV016, Etesevimab) and P2C-1F11 (BRII-196, Amubarvimab), IGHV3-53/3-66 public antibodies exhibit potent neutralizing activity against SARS-CoV-2 and provide effective protection against viral infection and disease progression in humans^1,12,13^. Nonetheless, most antibodies in this family have been escaped by VOCs, especially Omicron lineages that harbor extraordinarily high number of mutations in RBD^4,14-16^. Although a few antibodies, such as COV11, R40– 1G8, COVOX-222, P5-1C8, P5S-2B10, VacBB-551, BD55-1205 and BD56-1854, maintain broadly neutralizing activity across VOCs^8,17-21^, they are extremely rare given the prevalence of IGHV3-53/3-66 antibodies.

Understanding the characteristics of broadly neutralizing antibodies (bnAbs) against SARS-CoV-2 will guide the rational design of vaccines and antibody therapeutics with broad protection. In this study, we identified two convalescents of breakthrough infection with relatively high neutralizing titers against all tested viruses including BQ and XBB lineages. Among 50 spike-specific monoclonal antibodies (mAbs) cloned from their B cells, the top 6 neutralizing mAbs (KXD01-06) belong to IGHV3-53/3-66 public antibodies and exhibit broad neutralizing capacity. Deep mutational scanning reveals that they target highly conserved sites on RBD. Genetic and functional analysis indicates that the extent of somatic hypermutation is critical for their breadth. These findings provide rationale for the development of vaccines and antibody therapeutics based on IGHV3-53/3-66 public antibodies with broad and potent neutralizing activity.

## Materials and methods

### Human samples

Peripheral blood samples were collected from 11 donors (Table 1). Plasma and peripheral blood mononuclear cells (PBMCs) were separated from blood by Ficoll density gradient centrifugation. This study was approved by the Ethics Committee of Hefei Institutes of Physical Science, Chinese Academy of Sciences (Approval Number: YXLL-2023-47). All donors provided written informed consent for collection of information, analysis of plasma and PBMCs, and publication of data generated from their samples.

**Table 1.**
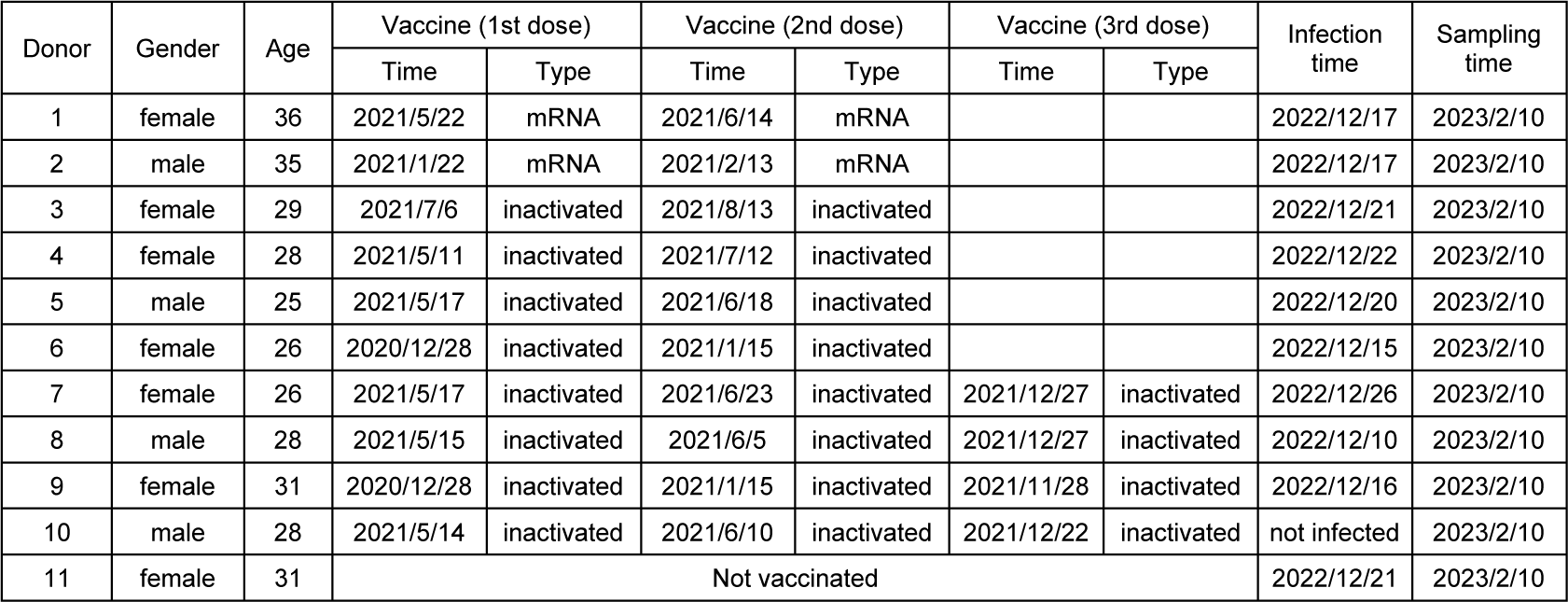
Information of the donors and samples.

| Donor | Gender | Age | Vaccine (1st dose) |  | Vaccine (2nd dose) |  | Vaccine (3rd dose) |  | Infection time | Sampling time |
| --- | --- | --- | --- | --- | --- | --- | --- | --- | --- | --- |
|  |  |  | Time | Type | Time | Type | Time | Type |  |  |
| 1 | female | 36 | 2021/5/22 | mRNA | 2021/6/14 | mRNA |  |  | 2022/12/17 | 2023/2/10 |
| 2 | male | 35 | 2021/1/22 | mRNA | 2021/2/13 | mRNA |  |  | 2022/12/17 | 2023/2/10 |
| 3 | female | 29 | 2021/7/6 | inactivated | 2021/8/13 | inactivated |  |  | 2022/12/21 | 2023/2/10 |
| 4 | female | 28 | 2021/5/11 | inactivated | 2021/7/12 | inactivated |  |  | 2022/12/22 | 2023/2/10 |
| 5 | male | 25 | 2021/5/17 | inactivated | 2021/6/18 | inactivated |  |  | 2022/12/20 | 2023/2/10 |
| 6 | female | 26 | 2020/12/28 | inactivated | 2021/1/15 | inactivated |  |  | 2022/12/15 | 2023/2/10 |
| 7 | female | 26 | 2021/5/17 | inactivated | 2021/6/23 | inactivated | 2021/12/27 | inactivated | 2022/12/26 | 2023/2/10 |
| 8 | male | 28 | 2021/5/15 | inactivated | 2021/6/5 | inactivated | 2021/12/27 | inactivated | 2022/12/10 | 2023/2/10 |
| 9 | female | 31 | 2020/12/28 | inactivated | 2021/1/15 | inactivated | 2021/11/28 | inactivated | 2022/12/16 | 2023/2/10 |
| 10 | male | 28 | 2021/5/14 | inactivated | 2021/6/10 | inactivated | 2021/12/22 | inactivated | not infected | 2023/2/10 |
| 11 | female | 31 | Not vaccinated |  |  |  |  |  | 2022/12/21 | 2023/2/10 |

### Cell lines

HEK-293T cells expressing human ACE2 (HEK-293T-hACE2) were kindly provided by Prof. Ji Wang at Sun Yat-Sen University. HEK-293T cells were from ATCC. HEK-293T and HEK-293T-hACE2 cells were cultured in DMEM with 10% FBS and 1% penicillin/streptomycin (pen/strep). FreeStyle 293 F cells (Thermo Fisher Scientific) were cultured in SMM 293-TII Expression Medium (Sino Biological Inc., M293TII). All cells were maintained in a 37°C incubator at 5% CO_2_.

### Protein expression and purification

The genes encoding the extracellular domain of SARS-CoV-2 spike including WT, Omicron BA.4/5, Omicron XBB.1.5 were constructed with a foldon trimerization motif and tandem strep tags at C-terminal. The genes encoding WT, BA.2.75, BA.4/5, BQ.1 and XBB.1.5 RBDs were constructed with a strep tag at C-terminal. Spike trimer and RBD were expressed in the FreeStyle 293 F cells and purified by BeaverBeads™ Strep-Tactin (BEAVER, 70808-250).

### ELISA

RBD or spike were coated onto 96-well ELISA plates (100ng/well) and incubated at 4°C overnight. After blocking with PBS containing 10% fetal bovine serum (FBS), 3– fold serially diluted plasma (starting at 1:100), mAbs (starting at 10μg/ml) or HEK293T supernatant were added to the wells and incubated at 37°C for 1hr. HRP–conjugated goat anti-human antibodies (Zen-bio, 550004; 1:5000 dilution) were added to the wells and incubated at 37°C for 1hr. TMB substrate (Sangon Biotech, E661007-0100) was added to the wells and incubated at room temperature for 5 mins. The reaction was stopped by TMB Stop Solution (Sangon Biotech, E661006-0500) and absorbance at 450nm were measured.

### Pseudovirus neutralization assay

To generate pseudoviruses, HEK-293T cells were transfected with psPAX2, pLenti-luciferase and spike-encoding plasmids using polyetherimide (PEI). Supernatant with pseudovirus was collected 48hrs after transfection. 3-fold serially diluted plasma (starting at 1:20), HEK293T supernatant (starting at 1:4.5), or mAbs (starting at 10μg/ml) were mixed with pseudoviruses at 37°C for 1hr. HEK293T-hACE2 cells (1.5X10^4^ per well) were added into the mixture and incubated at 37°C for 48hrs. Cells were lysed to measure luciferase activity (Bright-Lite Luciferase Assay System, DD1204-02 Vazyme Biotech Co.,Ltd). The percent of neutralization was determined by comparing with the virus control. The plasmids encoding spike of WT SARS-CoV-2, Delta, BA.1, BA.2, BA.3, BA.4/5, BF.7, BQ.1, XBB, XBB.1, XBB.1.5, XBB.1.16 and SARS-CoV-1 were kindly provided by Prof. Linqi Zhang at Tsinghua University. The plasmids encoding spike of BA.2.75 was kindly provided by Prof. Zezhong Liu at Fudan University. The plasmid encoding BA.4/5 spike with escape mutations were generated by PCR with primers containing mutations.

### Spike-specific Single B cell sorting and mAb cloning

PBMCs from Donor 1 and 2 were incubated with 200 nM SARS-CoV-2 spike for 30min at 4°C. After wash, they were stained with cell-surface antibodies: CD3-BV510 (BioLegend, 317331), CD19-PE/Cy7 (BioLegend, 302215), CD27-APC (BioLegend, 356409), CD38-APC/Cy7 (BioLegend, 356615), human IgM-AF700 (BioLegend, 314537), human IgD-perCP/Cy5.5 (BioLegend, 348207), anti-His-FITC (Proteintech, CL488-66005), anti-FLAG-PE (BioLegend, 637309) and DAPI. The stained cells were washed with FACS buffer (PBS containing 2% FBS) and resuspended in 500μl FACS buffer. Spike-specific single B cells were gated as DAPI^-^CD3^−^CD19^+^CD27^+^IgD^-^His^+^FLAG^+^ and sorted into 96-well PCR plates containing 4μl of lysis buffer (0.5XPBS, 0.1M DTT and RNase inhibitor) per well. After reverse transcription reaction, the heavy and light chain variable regions were amplified by nested PCR and cloned into expression vectors to produce full IgG1 antibodies^22^. The paired heavy– and light-chain were co-transfected into HEK-293T cells or FreeStyle 293 F cells. Antibodies were purified with Protein A magnetic beads (GenScript, L00273). Expression vectors for CoV-2196, CoV-2130, P2C-1F11, P5-2H11, P5S-2B10 were kingly provided by Prof. Linqi Zhang at Tsinghua University. The variable regions of BD55-1205 and BD56-1854 were synthesized and cloned into expression vectors.

### Competition ELISA

WT RBD was coated onto 96-well ELISA plates (100ng/well) and incubated at 4°C overnight. After blocking with PBS containing 10% FBS, biotinylated ACE2, CoV-2196, CoV-2130 and P2C-1F11 were mixed with competing mAbs at 1:50 molar ratio. The mixtures were added to the wells and incubated at 37°C for 1hr. Streptavidin-HRP (GenScript, M00091; 1:5000 dilution) was added to the wells and incubated at 37°C for 1hr. TMB substrate (Sangon Biotech, E661007-0100) was added to the wells and incubated at room temperature for 5-10 mins. The reaction was stopped by TMB Stop Solution (Sangon Biotech, E661006-0500) and absorbance at 450nm were measured. The percentages of signal decrease caused by competing mAbs were calculated.

### Deep mutational scanning

Yeast libraries displaying BA.4/5 RBD mutants were kindly provided by Prof. Yunlong Cao at Peking University. Three rounds of FACS were performed to enrich RBD mutants losing binding to mAb but maintaining binding to ACE2. In the first round, yeasts were stained with ACE2 and ACE2-positive yeasts were sorted and expanded. In the second round, yeasts were stained with mAbs and mAb-negative yeasts were sorted and expanded. In the third round, yeasts were stained with ACE2 and mAbs simultaneously. ACE2-postive but mAb-negative yeasts were sorted. With the yeasts post the third sorting as template, PCR was performed to amplify RBD fragment from the plasmid. The PCR products were cloned to T vector and sequenced by Sanger-sequencing.

## Data analysis

Sequences of antibody heavy and light chain variable regions from single B cells were analyzed with IgBlast (https://www.ncbi.nlm.nih.gov/igblast/). Sequence alignment was performed either by MEGA or by the Muscle v5 algorithm. Sequence logos displaying mutation profiles were created with the Pandas, Bio, Matplotlib, Seaborn, and Logomaker packages in Python 3.8.16 environment. FACS data were analyzed with Flowjo. ELISA and neutralizing data were analyzed and plotted with Graphpad Prism 8.

## Results

### Identification of individuals with broad neutralizing activity

To characterize bnAbs elicited by infection or vaccination, we collected peripheral blood samples from 11 donors (Table 1). Except Donor 11 who was not vaccinated with any SARS-CoV-2 vaccine, all the other donors were vaccinated with either two doses of mRNA vaccine (Donor 1 and 2), or two doses of inactivated vaccine (Donor 3-6), or three doses of inactivated vaccine (Donor 7-10). During the wave of infection in December 2022, all donors except Donor 10 were infected with SARS-CoV-2, probably BA.5 or BF.7 variants as they were main circulating strains at that time^23^.

We measured plasma binding antibody titers against prototype (wild-type, WT), BA.4/5 and XBB.1.5 spike by ELISA (Figure 1A, B). The antibody titers vary in a wide range with highest titers from Donor 2 and lowest titers from Donor 10 and 11. Moreover, antibody titers against WT, BA.4/5 and XBB.1.5 spike gradually decrease, which correlate with the number of mutations on these spikes. We further measured plasma neutralizing antibody titers against 14 pseudoviruses including WT, Delta, BA.1, BA.2, BA.3, BA.4/5, BA.2.75, BF.7, BQ.1, XBB, XBB.1, XBB.1.5, XBB.1.16 and SARS-

**Figure 1.**
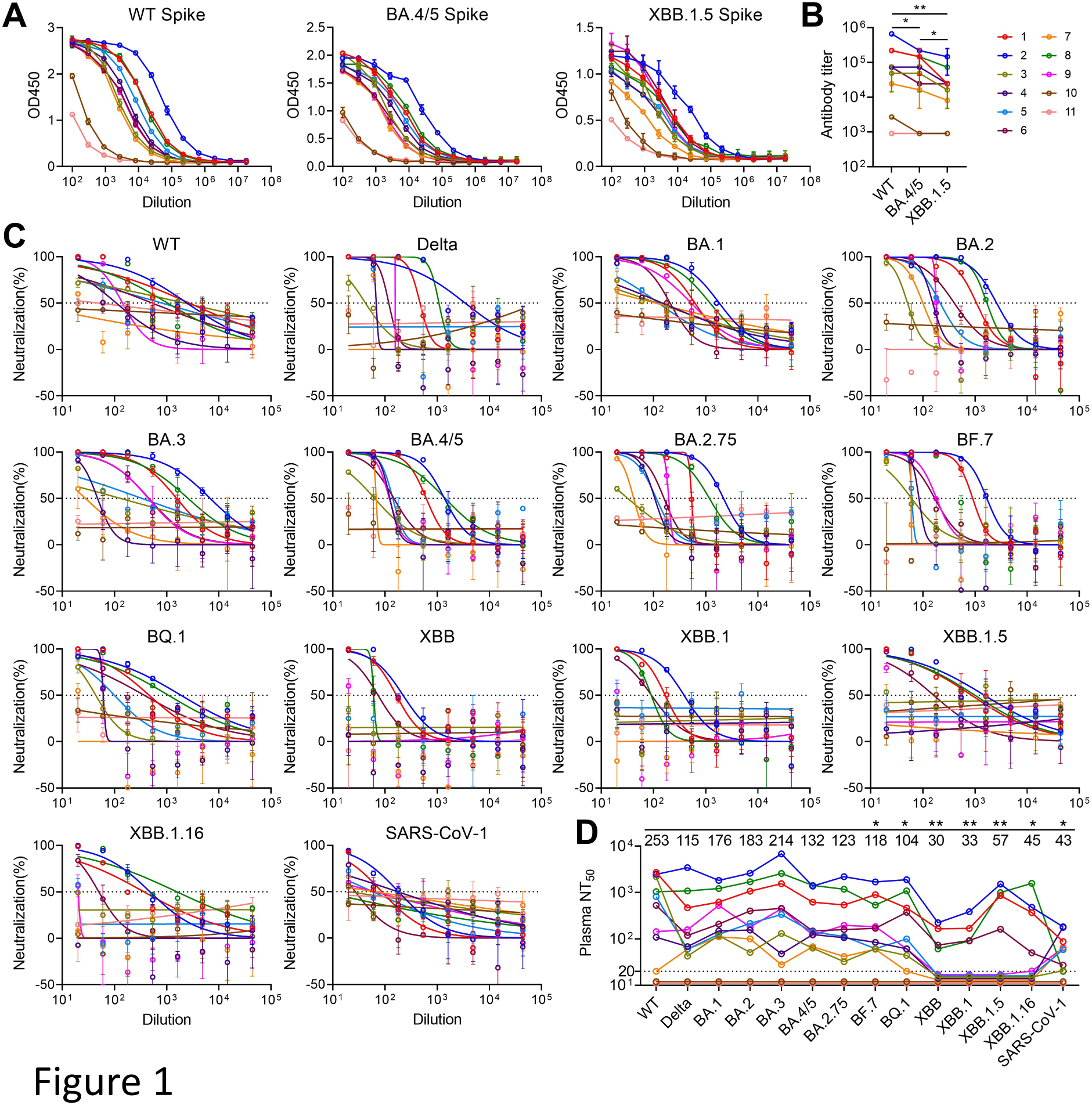
Characterization of plasma from SARS-CoV-2 convalescents or vacinees. (**A**) Measurement of antibody titers against WT, BA.4/5 and XBB.1.5 spikes by ELISA. Data are represented as the mean ± SD. (**B**) Summary of antibody titers. Statistical analysis was performed by two-tailed Wilcoxon matched-pairs signed rank test. *P < 0.05, **P < 0.01. (**C**) Measurement of neutralizing antibody titers against a panel of pseudovirues. Data are represented as the mean ± SD. (**D**) Summary of neutralizing antibody titers. For plasma with no neutralizing activity at 20-fold dilution, a number between 10-20 is given as titer to separate the curves. The numbers on top are Geometric mean titers against the viruses. The titers against WT are compared with titers against other viruses. Statistical analysis was performed by two-tailed Wilcoxon matched-pairs signed rank test. *P < 0.05, **P < 0.01.

CoV-1 (Figure 1C, D). Donor 10 and 11 show little neutralizing activity, consistent with their low binding antibody titers. In contrast, Donor 1, 2, 6 and 8 exhibit broad neutralizing activity against all tested viruses. Donor 3, 4, 5, 7 and 9 have neutralizing titers against viruses other than XBB lineages. The geometric mean titers against each virus further confirm that XBB lineages are most resistant to neutralization, which is consistent with other studies^5–8^.

### Isolation of mAbs from two convalescents

As Donor 1 and 2 have higher neutralizing titers against XBB and XBB.1 than other donors, we chose their blood cell samples to isolate mAbs. With WT spike as bait, we sorted single antigen-specific memory B cells and plasmablasts (Figure S1A). From those cells, we cloned 72 mAbs by single cell RT-PCR (Figure 2A, B, Figure S1B). By screening supernatant of 293T cells transfected with antibody-expressing vectors, we identified 50 mAbs positive for WT spike, among which 25 mAbs are also positive for WT RBD (Figure 2A, B). We further tested the neutralizing activity of spike-positive supernatant against WT pseudovirus. In sum, we identified 6 mAbs (KXD01-06) with potent neutralizing activity and 3 mAbs (KXD07-09) with moderate neutralizing activity (Figure 2B). Notably, KXD01-06 are all encoded by IGHV3-53/3-66, with either short CDRH3 ranging from 10 to 12 amino acids or long CDRH3 of 21 amino acids. Moreover, light chain V genes of these mAbs, including IGKV1-9, IGKV1-33, IGKV3-11 and IGKV3-15, have been frequently reported to pair with IGHV3-53/3-66 (Figure 2C). These data suggest that the top neutralizing antibodies from these two convalescents belong to previously defined IGHV3-53/3-66 public antibodies.

**Figure 2.**
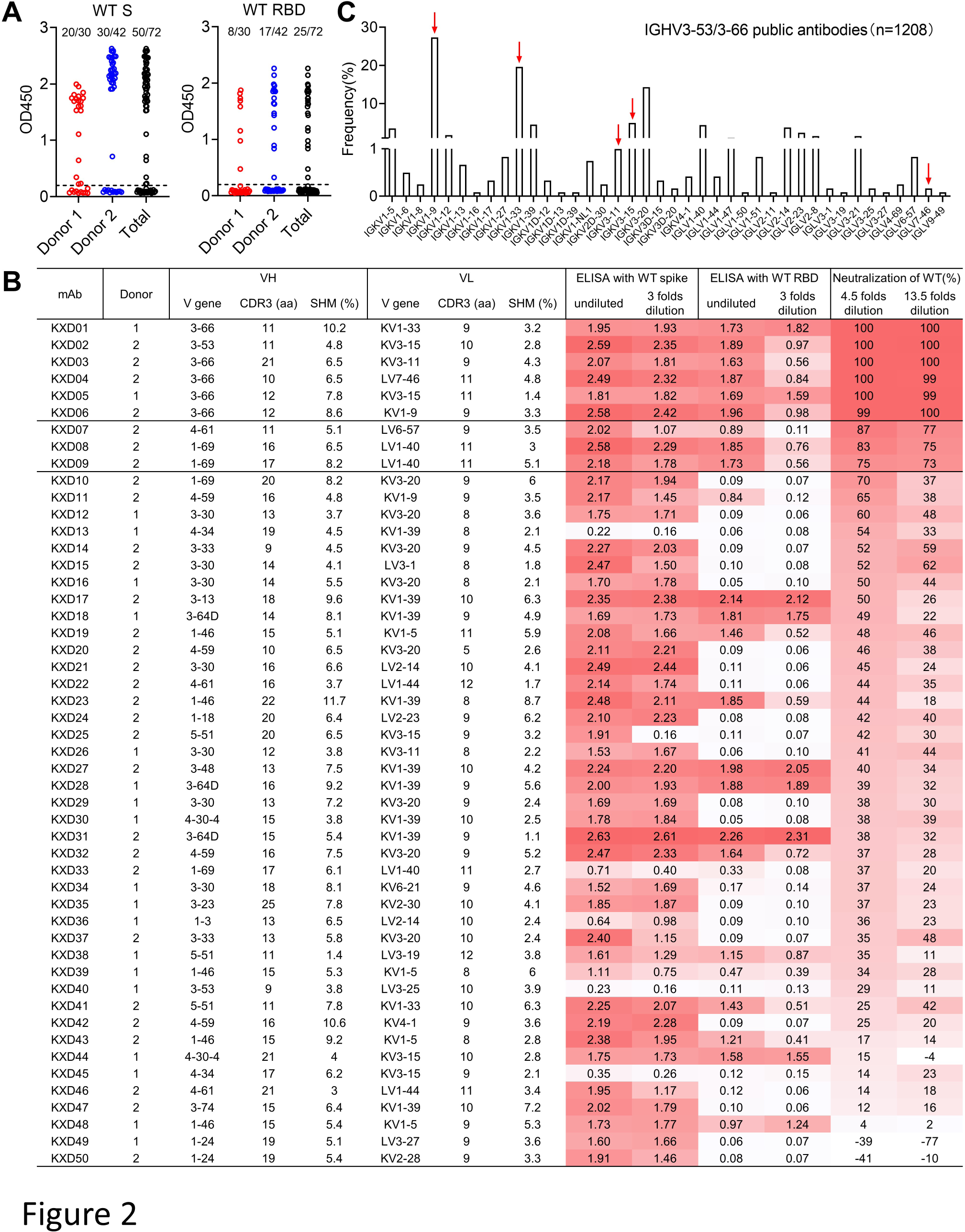
Isolation and characterization of mAbs from Donor 1 and 2. (A) Summary of mAb screening by ELISA. Undiluted supernatant from 293T cells transfected with antibody-expressing vectors was tested against WT spike and RBD. OD450 was measured and positive clones were identified with 0.2 as cut-off. (B) Summary of 50 spike positive mAbs. The mAbs are ordered based on the neutralizing activity of 293T supernatant. OD450s and neutralization percentages are color-coded, with darker red indicates higher values. (C) Frequency of light chain V gene usage among 1208 IGHV3-53/3-66 public antibodies. The red arrows indicate V genes used by KXD01-06.

### Characterization of KXD01-09

To further characterize those neutralizing mAbs, we produced KXD01-09 together with CoV-2196, CoV-2130, P2C-1F11, P5-2H11, P5S-2B10, BD55-1205 and BD56-1854. CoV-2196, CoV-2130 and P2C-1F11 are the prototypes of Tixagevimab, Cilgavimab and Amubarvimab, which was approved for emergency use^1^. In addition to P2C-1F11, P5-2H11, P5S-2B10, BD55-1205 and BD56-1854 are also IGHV3-53/3-66 public antibodies^8, 20^. Moreover, P5S-2B10, BD55-1205 and BD56-1854 have broad neutralizing activity across VOCs.

We performed competition ELISA with biotin-labeled ACE2, CoV-2196, CoV-2130 and P2C-1F11 (Figure 3). As expected, KXD01-06 block the binding of RBD with ACE2, CoV-2196 and P2C-1F11, further confirming that they target similar epitope as other IGHV3-53/3-66 public antibodies and neutralize SARS-CoV-2 by ACE2 blocking. Although KXD07-09 compete with CoV-2130, they do not compete with ACE2 like CoV-2130. Therefore, the epitopes of KXD07-09 remain elusive.

**Figure 3.**
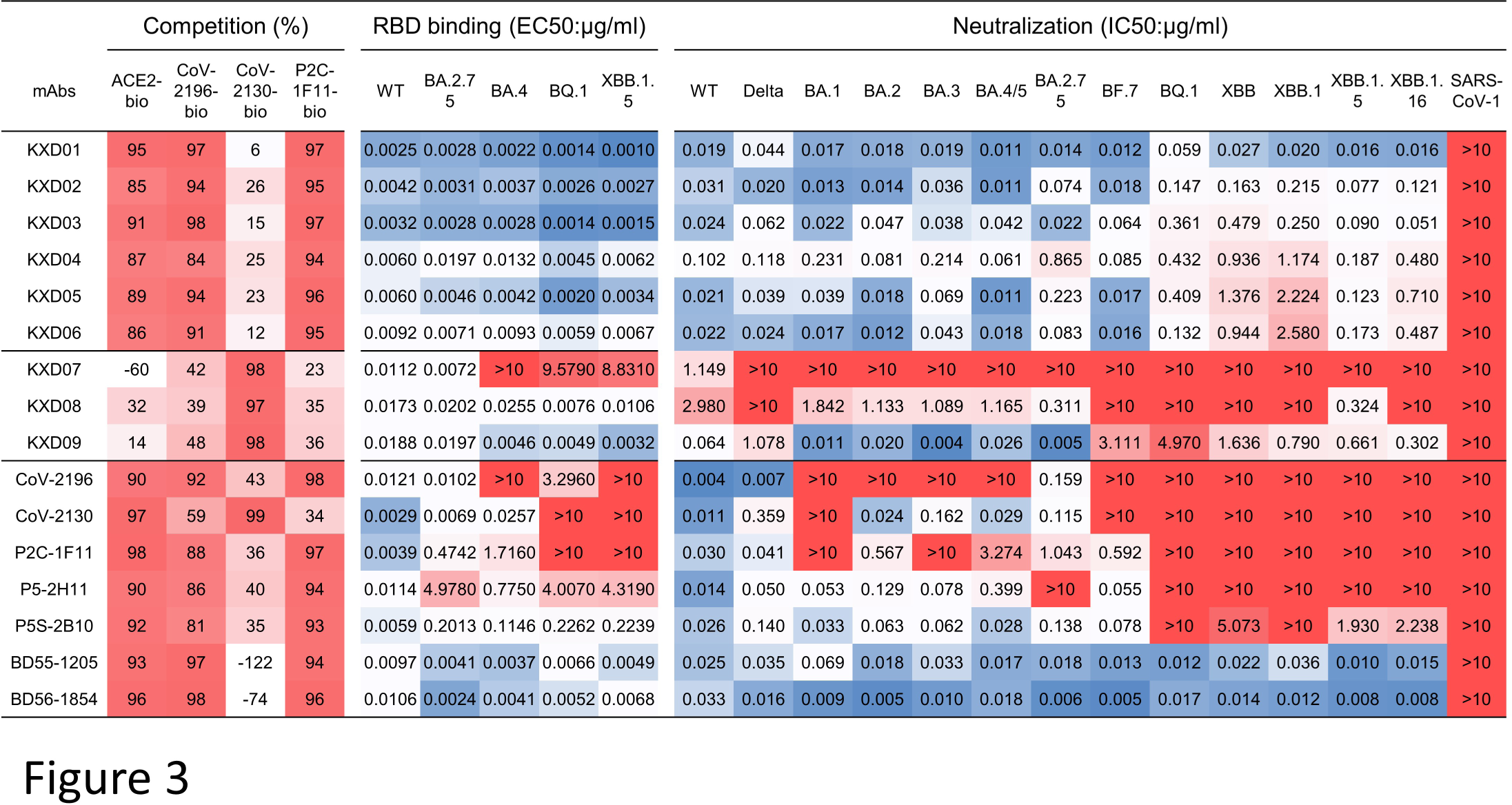
Comparison of epitope, binding activity and neutralizing activity among KXD01-09 and reported antibodies. Epitopes were mapped by competition ELISA. ACE2, CoV-2196, CoV-2130 and P2C-1F11 were labeled with biotin. The percentages of their binding to RBD competed by KXD01-09 and reported antibodies were measured by ELISA. The values are color-coded with darker red indicates more competition. The binding activity against WT and variant RBDs was measured by ELISA. The neutralizing activity was measured by pseudoviruses. The highest antibody concentration used to determine EC50 and IC50 is 10μg/ml. EC50s and IC50s were calculated by least squares fit. The values of EC50s and IC50s are color-coded with dark blue indicates lower values and dark red indicates higher values.

We also measured the binding activity of these mAbs against WT and variant RBDs by ELISA (Figure 3, Figure S2A). All KXD mAbs except KXD08 maintain binding activity to all tested RBDs. Among the published mAbs, BD 55-1205 and BD56-1854 show consistent binding to all RBDs, whereas the other mAbs lose binding to variant RBDs with different degrees.

We also tested the neutralizing activity of these mAbs against the same panel of pseudovirues used for plasma samples (Figure 3, Figure S2B). Surprisingly, KXD01-06 are able to neutralize all SARS-CoV-2 viruses. The IC50s of the best mAb KXD01 range from 0.011∼0.059μg/ml with geometric mean of 0.020μg/ml, whereas KXD02-06 show 2 to 100 folds reduction in neutralization against BA.2.75, BQ.1 and XBB lineages compared with WT. KXD09 is another bnAb with reduced neutralizing potency against multiple variants including Delta, BF.7, BQ.1 and XBB lineages. KXD07 and KXD08 have limited neutralizing breath and potency. Among the published mAbs, BD55-1205 and BD56-1854 show broad and potent neutralizing antibodies against SARS-CoV-2 viruses, while the other antibodies are largely escape by variants, particularly BQ.1 and XBB lineages. Taken together, we identified 6 IGHV3-53/3-66 public antibodies with broad neutralizing capacity against SARS-CoV-2.

### Mapping escape mutations of KXD01-06

To explore the molecular basis for broad neutralization and neutralization escape, we performed deep mutational scanning (DMS) to map escape mutations of KXD01-06 (Figure 4A, Figure S3). We first filtered out RBD mutants losing binding to ACE2 from two independently-constructed mutant libraries based on BA.4/5 RBD. Then we sorted RBD mutants with reduced binding to antibodies. In previously reported method^24, 25^, mutants post second sorting are directly processed to sequencing. Here we performed an additional round of sorting to further remove mutants maintaining binding to antibodies or losing binding to ACE2. Sequence analysis of mutants post third sorting shows that escape mutations of KXD01-06 are limited to residues including D420, Y421, L455, F456, N460, A475 and N487. In addition, the escape mutations of each antibody display a different pattern regarding to frequency, implying that there are minor differences in their interactions with RBD (Figure 4B).

**Figure 4.**
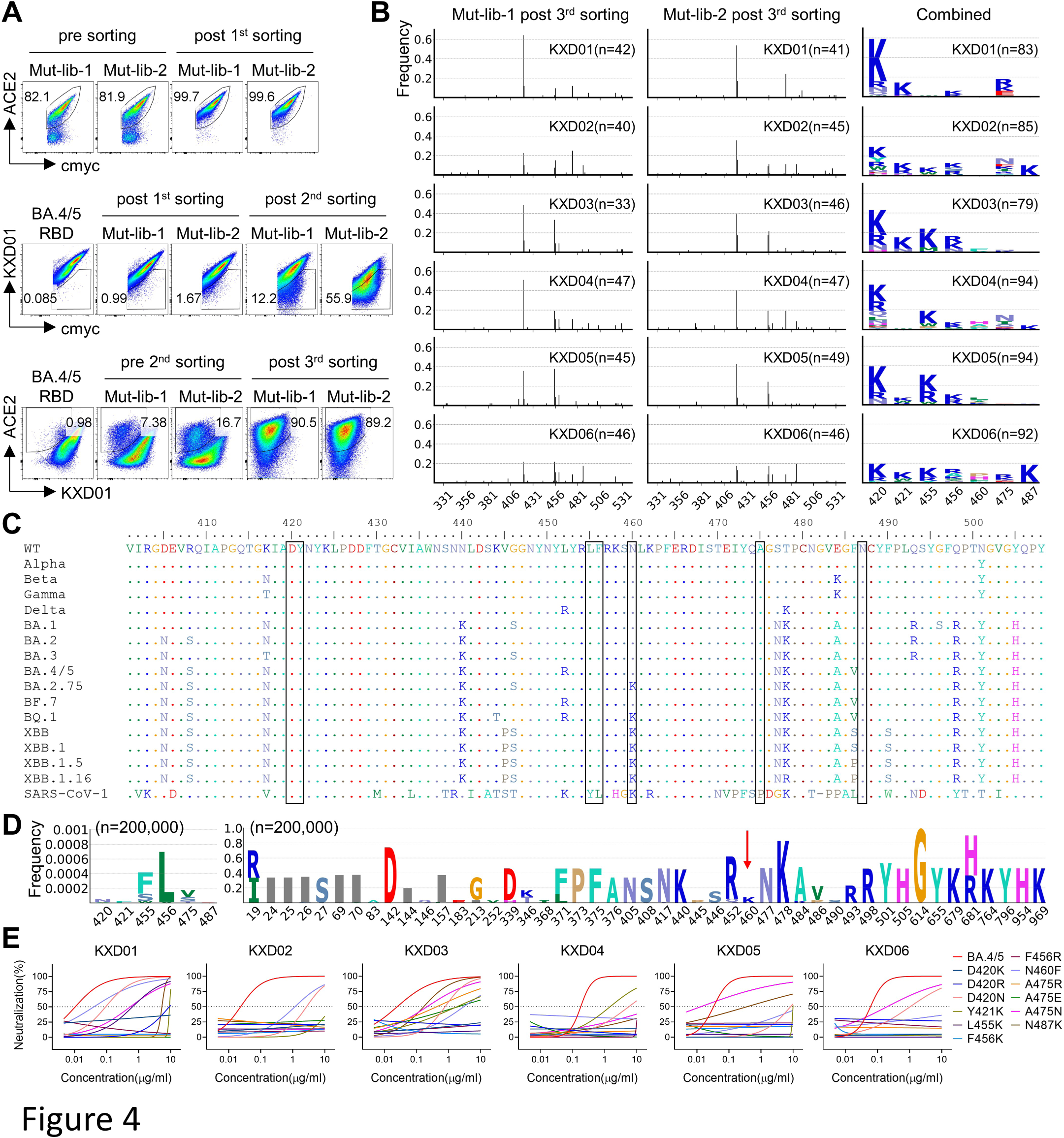
Mapping of escape mutation. (A) FACS plots illustrating the process of library screening. In the first sorting, ACE-2-positive mutants were sorted. In the second sorting, mAb-negative mutants were sorted. In the third sorting, ACE2-positive but mAb-negative mutants were sorted. (B) Sequence analysis of mutants post the third sorting. The sequences were aligned with BA.4/5 RBD and then the mutations were identified. The left two columns represent mutation frequency across RBD (331-531). The right column represents both mutations and frequencies on 420, 421, 455, 456, 460, 475 and 487. (C) Mutation profile of representative spike sequences. (D) Mutation profile of 200,000 SARS-CoV-2 spike sequences. The red arrow indicates position 460. (E) Neutralizing activity of KXD01-06 against BA.4/5 with escape mutations. Data are represented as non-linear fit curves calculated by least squares fit.

We analyzed the variations on the 7 residues among sequences of WT SARS-CoV-2, representative VOCs and SARS-CoV-1 (Figure 4C). No mutations are found on D420, Y421 and N487 and mutations are only found in SARS-CoV-1 on L455, F456 and A475. In contrast, N460K is a common mutation shared by BA.2.75, BQ.1, XBB lineages and SARS-CoV-1. We also randomly collected a total of 200,000 spike sequences and analyzed the mutation frequency on each residue (Figure 4D). The mutation frequencies on D420, Y421, L455, F456, A475 and N487 are nearly 0, with 5X10^-5^, 2.5X10^-5^, 4.2X10^-4^, 7.0X10^-4^, 1.8X10^-4^ and 5X10^-6^ respectively, whereas the mutation frequency on N460 is 0.077 because of N460K. These results suggest that all the residues except N460 are highly conserved among SARS-CoV-2 variants.

To confirm the effects of mutations on those residues to neutralization evasion, we measured the neutralizing activity of KXD01-06 against BA.4/5 pseudoviruses with escape mutations (Figure 4E). Overall, KXD01-06 are largely or completely escaped by viruses carrying mutations identified by DMS. It is noteworthy that KXD01 maintains relatively potent neutralizing activity against virus with N460F, consistent with above neutralization profile of KXD01 against variants with N460K such as BA.2.75, BQ.1, XBB lineages. Taken together, these results demonstrate that KXD01-06 target highly conserved epitopes and they can be escaped by future variants with mutations on residues including D420, Y421, L455, F456, N460, A475 and N487.

### Genetic basis for broad neutralizing activity of KXD01-06

To illuminate the formation of broad neutralizing activity, we move on to characterize the genetic features of KXD01-06 compared to other IGHV3-53/3-66 public antibodies. We collected 1208 published antibodies and analyzed their mutations. As previously reported, IGHV3-53/3-66 public antibodies generally have limited mutations on IGHV3-53/3-66, with 62.1% antibodies have 0-5 mutations and only 8.7% antibodies have more than 10 mutations (Figure 5A). KXD01-06 have more mutation than most IGHV3-53/3-66 antibodies, with 18, 6, 11, 10, 9 and 12 mutations respectively (Mutations probably introduced by PCR primers at the beginning of IGHV3-53/3-66 are not counted) (Figure 5B, C). We also summarized the mutation profile of published IGHV3-53/3-66 antibodies, which shows that they share common mutations such as Y57F, F27L, F27I, T28I, S31R, S35N, S35I and V50L (Figure 5C). Referring to this profile, the abundant mutations accumulated by KXD01-06 are rather common instead of rare or unique.

**Figure 5.**
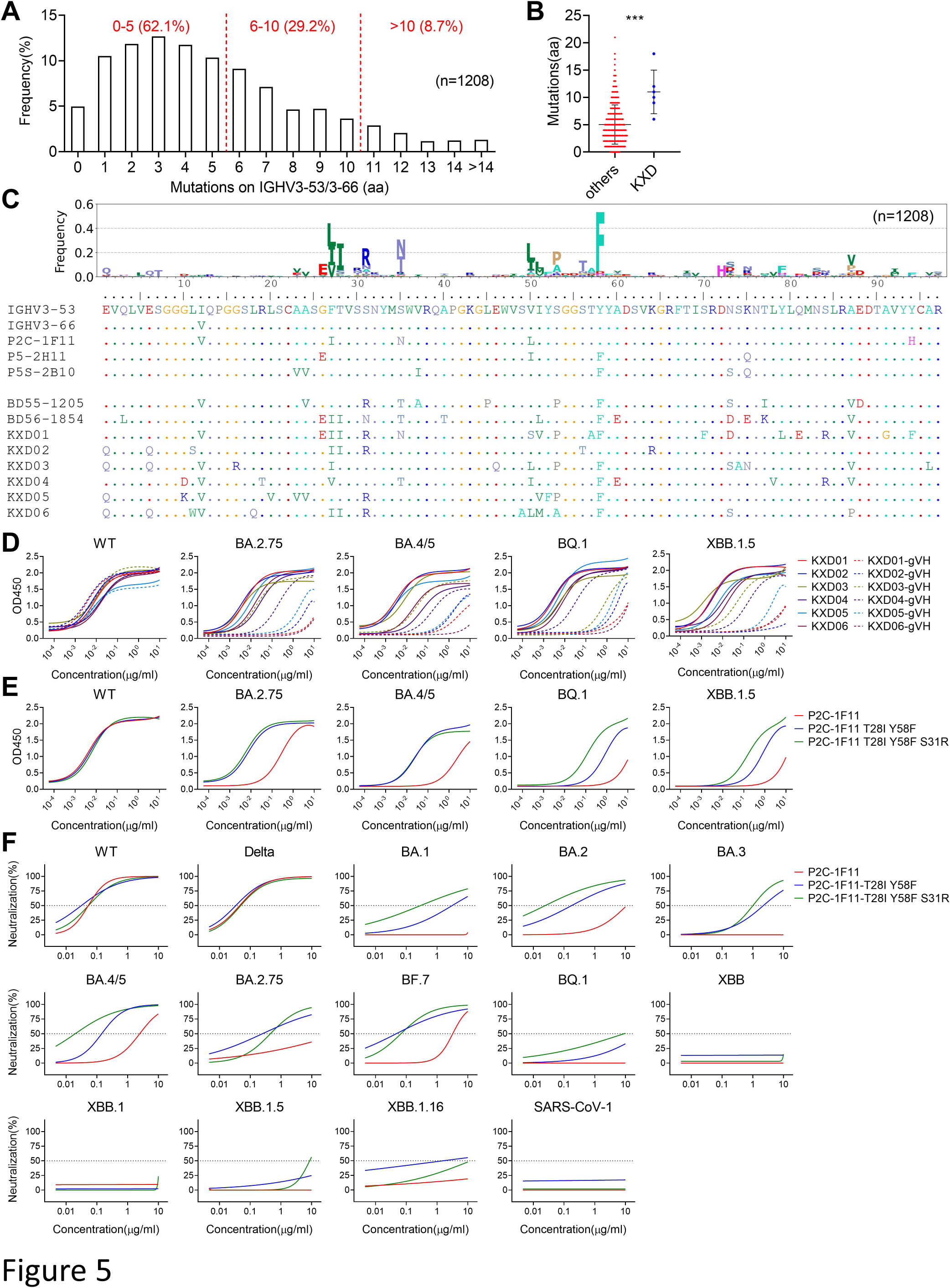
Effects of somatic hypermutation on neutralization breadth. (A) Summary of mutations on IGHV3-53/3-66 accumulated by 1208 IGHV3-53/3-66 public antibodies against SARS-CoV-2. (B) Mutation levels of KXD01-06 and 1208 reported antibodies. (C) Mutation profiles of KXD01-06 with other reported antibodies. (D) Binding activity of KXD01-06 and their germline-VH versions measured by ELISA. (E) Binding activity of P2C-1F11 and P2C-1F11 with additional mutations measured by ELISA. (F) Neutralizing activity of P2C-1F11 and P2C-1F11 with additional mutations against a panel of pseudoviruses. The binding and neutralizing data are represented as non-linear fit curves calculated by least squares fit.

To investigate the function of those mutations, we reverted the IGHV region of KXD01-06 to IGHV3-53/3-66 germline sequences. Then we compared the binding activity of mature antibodies and germline antibodies to WT and variant RBDs (Figure 5D). Although each pair of antibodies bind WT RBD with similar affinities, the germline versions partially or almost completely lose binding to BA.2.75, BA.4/5, BQ.1 and XBB.1.5 RBDs. Oppositely, we introduced common mutations including T28I, S31R and Y58F to IGHV region of P2C-1F11 as it lacks these mutations. Consistent with above results, these mutations dramatically increase the binding activity of P2C-1F11 to variant RBDs while not to WT RBD (Figure 5E). Furthermore, these mutations substantially improve the neutralizing activity of P2C-1F11 against BA.1, BA.2, BA.2.75, BA.3, BA.4/5 and BF.7, although BQ.1 and XBB lineages are still resistant to neutralization. Taken together, these results elucidate that the extent of somatic hypermutation is critical for the neutralization breadth of IGHV3-53/3-66 public antibodies.

## Discussion

In this study, we mainly characterized the antibody response in two convalescents recovered from breakthrough infection probably caused by SARS-CoV-2 BA.5 or BF.7. The top 6 neutralizing mAbs (KXD01-06) among a total of 50 spike-specific mAbs cloned from their B cells are encoded by IGHV3-53/3-66, which once again confirms the dominance of IGHV3-53/3-66 antibodies in neutralizing antibodies against SARS-CoV-2. It is surprising that KXD01-06 have broad neutralizing activity across VOCs including BQ and XBB lineages, as most IGHV3-53/3-66 antibodies identified previously have been escaped by VOCs, particularly Omicron variants^4,14-16^. Several groups reported IGHV3-53/3-66 antibodies with broad neutralizing activity against Omicron variants^17–20^, whereas it remains to be tested whether those antibodies are escaped by BQ and XBB lineages. Recently, Cao et al. comprehensively measured the neutralizing activity of more than 3000 mAbs against a panel of SARS-CoV-2 pseudoviruses including D614G, BA.1, BA.2, BA.2.75, BA.5, BQ.1.1 and XBB. According to this dataset, 32 out of 181 IGHV3-53/3-66 antibodies broadly neutralize all tested viruses, with 3 from SARS convalescents, 2 from WT vaccinees, 4 from WT convalescents, 7 from BA.1 convalescents, 9 from BA.2 convalescents and 7 from BA.5 convalescents^8^. Considering those mAbs were identified by high-throughput single-cell sequencing of hundreds of thousands of B cells, the frequency of broadly neutralizing IGHV3-53/3-66 antibodies in those convalescents and vaccinees are much lower than these two convalescents in this study.

The development of bnAbs against highly variable pathogens is a key question for immunology and vaccinology. Currently, the development of HIV-1 bnAbs is most studied and a co-evolution model is proposed based on longitudinal analysis of the race between viruses and B cell lineages^26^. According to this model, envelop glycoprotein (Env) from transmitted/founder (T/F) virus leads the rare B cell precursors of bnAbs to undergo clonal expansion and somatic hypermutation. Subsequently, B cell lineages with desired mutations are selected by Envs from variants of T/F virus for further diversification. After iterative selections, B cell lineages gradually accumulate high levels of somatic hypermutations and eventually mature to produce bnAbs. Overall, the development of HIV-1 bnAbs is a rare event due to the following roadblocks. First, the precursor B cells of HIV-1 bnAbs are extremely rare in human B cell repertoire^27^. Second, those precursor B cells generally show poor binding activity with most Envs^28, 29^. Third, HIV-1 bnAbs have 10%∼30% mutations and some mutations are intrinsically improbable^30, 31^. Compared with HIV-1 bnAbs, there are no such roadblocks in the development of broadly neutralizing IGHV3-53/3-66 antibodies against SARS-CoV-2. As the dominant antibodies targeting RBD, IGHV3-53/3-66 antibodies are prevalent in human B cell repertoire^32, 33^. Moreover, IGHV3-53/3-66 germline antibodies already have moderate to high affinities to SARS-CoV-2^34, 35^. In addition, our study shows that the mutation level of broadly neutralizing IGHV3-53/3-66 antibodies is much lower than HIV-1 bnAbs and those mutations are commonly found in IGHV3-53/3-66 antibodies. These findings suggest that IGHV3-53/3-66 antibodies are promising targets for vaccines aiming to elicit bnAbs against SARS-CoV-2.

Regarding to mutations accumulated during antibody affinity maturation, we show that the mutations on IGHV3-53/3-66 substantially enhance the binding activity of KXD01-06 to variant RBDs instead of WT RBD, and thus we speculate that those mutations are selected by variant RBDs during breakthrough infection, which is consistent with co-evolution model of HIV-1 bnAbs. However, IGHV3-35/3-66 antibodies with broad neutralizing activity are also identified in WT vaccinees and WT convalescents as mentioned above, suggesting that variants are not essential for the development of neutralizing breadth across VOCs. Therefore, more studies are required to understand the development of IGHV3-53/3-66 antibodies with broad neutralizing activity.

According to the classification of RBD-specific antibodies^9,36,37^ (ref), IGHV3-53/3-66 antibodies mainly fall into two groups: RBD class 1 (RBS-A; RBD-2a) and RBD class 2 (RBS-B; RBD-2b). IGHV3-53/3-66 antibodies in RBD class 1 have short CDRH3 and bind to RBM using germline-encoded NY and SGGS motifs in CDRH1 and CDRH2. They only bind to RBD in the up conformation and neutralize SARS-CoV-2 by ACE2 blocking. In contrast, IGHV3-53/3-66 antibodies in RBD class 2 have long CDRH3 and can bind to RBD in both up and down conformation. They mainly contact with the RBD ridge and its nearby regions, and also neutralize SARS-CoV-2 by ACE2 blocking. According to this classification, KXD03, which has a long CDRH3 with 21 amino acids, is supposed to target different epitopes with the other 5 mAbs. Indeed, KXD01-06 share similar escape map with P2C-1F11, which is a representative of RBD class 1 antibody with escape mutations distributed on D420, Y421, L455, F456, N460, P463, Y473, A475 and N487^25^. Therefore, IGHV3-53/3-66 antibodies with long CDRH3 can target similar epitopes as IGHV3-53/3-66 antibodies with short CDRH3.

We acknowledge there are several potential limitations of this study. First, we only identified 50 spike-specific mAbs, which may be not enough to represent the antibody response in the two individuals. Second, we did not collect blood samples before the breakthrough infection. So we are not able to compare IGHV3-53/3-66 antibodies generated before and after the breakthrough infection, which is helpful to understand the development of broadly neutralizing IGHV3-53/3-66 antibodies. Finally, the profiles of neutralization and escape mutations suggests that there are some minor differences in the interactions of KCD01-06 with spike. In future studies, structure analysis can be performed to further elucidate these differences.

The rapid emergence of SARS-CoV-2 VOCs highlights the urgent need to develop vaccines and antibody therapeutics with broad protection against current and future SARS-CoV-2 variants. This study demonstrates that IGHV3-53/3-66 public antibodies have enormous potential to develop broad and potent neutralizing activity through antibody affinity maturation, which provides rationale for novel vaccines and antibody therapeutics based on this class of antibodies.

## Supplemental information

Supplemental information includes three figures.

## Declaration of interests

L.J., T.Z., L.L. and X.C. are inventors on a patent application for antibodies including KXD01-06. The other authors declare that they have no competing interests.

## Acknowledgments

The authors thank all donors for providing the blood samples. The authors thank Prof. Linqi Zhang at Tsinghua University, Prof. Ji Wang at at Sun Yat-Sen University, Prof. Yunlong Cao at Peking University and Prof. Zezhong Liu at Fudan University for providing key materials for this study. The authors thank Zexuan Zheng at University of Science and Technology of China for help with yeast sorting. This work was supported by the National Key Plan for Scientific Research and Development of China (Grant No. 2022YFC2305800) and the National Natural Science Foundation of China (Grant No. 32200765).

## Figure legends

**Figure S1.** Isolation and characterization of mAbs from Donor 1 and 2. (A) FACS plots representing gating strategies for spike-specific single B cell sorting. (B) Summary of 22 spike-negative mAbs.

**Figure S2.** Measurement of binding activity and neutralizing activity of KXD01-09 and reported antibodies. (A) Binding activity against WT and variant RBDs measured by ELISA. (B) Neutralizing activity against a panel of pseudoviruse. The data are represented as non-linear fit curves calculated by least squares fit.

**Figure S3.** FACS plots of the second and the third sorting of two RBD mutant libraries by KXD01-06.

