## Supplementary material for "Breakthrough infection elicits hypermutated IGHV3-53/3-66 public antibodies with broad and potent neutralizing activity against SARS-CoV-2 variants including BQ and XBB lineages": suplemental figures

A

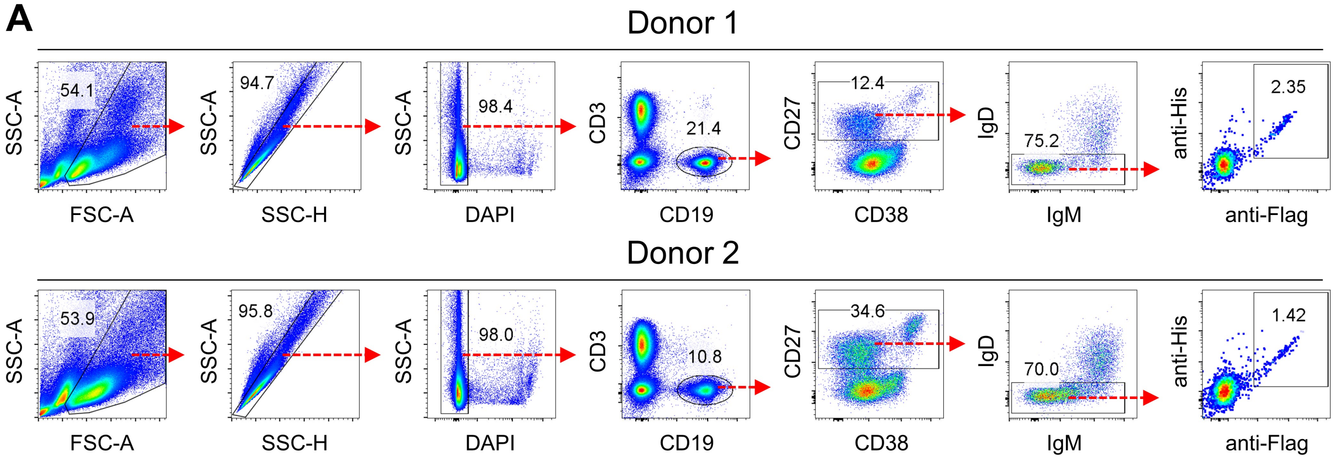


B

| mAb | Donor | VH | | | VL | | | ELISA with WT spike | | ELISA with WT RBD | |
|---|---|---|---|---|---|---|---|---|---|---|---|
| | | V gene | CDR3 (aa) | SHM (%) | V gene | CDR3 (aa) | SHM (%) | undiluted | 3 folds dilution | undiluted | 3 folds dilution |
| KXD51 | 1 | 1-18 | 22 | 3.7 | LV2-11 | 10 | 4.9 | 0.15 | 0.14 | 0.07 | 0.13 |
| KXD52 | 2 | 1-69 | 17 | 12.8 | KV1-39 | 9 | 8.3 | 0.13 | 0.12 | 0.17 | 0.16 |
| KXD53 | 2 | 3-48 | 13 | 7.8 | KV3-15 | 10 | 4.5 | 0.12 | 0.08 | 0.12 | 0.07 |
| KXD54 | 1 | 1-69 | 16 | 23 | KV1-39 | 9 | 8.1 | 0.12 | 0.16 | 0.05 | 0.08 |
| KXD55 | 2 | 7-4-1 | 10 | 3.4 | LV6-57 | 9 | 2.8 | 0.11 | 0.08 | 0.12 | 0.08 |
| KXD56 | 2 | 5-51 | 20 | 6.3 | KV3-20 | 10 | 4.9 | 0.10 | 0.10 | 0.09 | 0.12 |
| KXD57 | 1 | 3-53 | 10 | 3.4 | LV1-40 | 12 | 2.4 | 0.10 | 0.11 | 0.09 | 0.08 |
| KXD58 | 2 | 3-7 | 11 | 7.1 | KV2-30 | 9 | 2 | 0.09 | 0.10 | 0.10 | 0.13 |
| KXD59 | 2 | 1-69 | 17 | 15.9 | KV1-39 | 9 | 10.6 | 0.09 | 0.09 | 0.09 | 0.12 |
| KXD60 | 2 | 1-46 | 18 | 11.3 | LV2-23 | 11 | 6.5 | 0.09 | 0.10 | 0.09 | 0.13 |
| KXD61 | 1 | 3-53 | 9 | 3.6 | LV6-57 | 9 | 6.5 | 0.08 | 0.10 | 0.11 | 0.09 |
| KXD62 | 2 | 1-46 | 19 | 0.3 | KV1-39 | 9 | 0.7 | 0.08 | 0.08 | 0.08 | 0.07 |
| KXD63 | 2 | 3-23 | 14 | 0.7 | KV1-5 | 9 | 2.6 | 0.08 | 0.06 | 0.08 | 0.07 |
| KXD64 | 2 | 4-61 | 18 | 12 | KV3-15 | 4 | 4.7 | 0.08 | 0.09 | 0.08 | 0.08 |
| KXD65 | 1 | 4-34 | 19 | 8.9 | KV1-33 | 9 | 4.7 | 0.08 | 0.11 | 0.07 | 0.08 |
| KXD66 | 2 | 4-61 | 11 | 10 | LV3-1 | 9 | 5.8 | 0.08 | 0.07 | 0.10 | 0.08 |
| KXD67 | 1 | 3-53 | 9 | 4.1 | KV3-15 | 9 | 1 | 0.07 | 0.13 | 0.05 | 0.10 |
| KXD68 | 2 | 5-51 | 16 | 1 | LV1-40 | 10 | 0.7 | 0.07 | 0.06 | 0.08 | 0.06 |
| KXD69 | 1 | 3-33 | 17 | 7.1 | LV1-44 | 12 | 2.1 | 0.07 | 0.12 | 0.06 | 0.10 |
| KXD70 | 1 | 1-46 | 11 | 25.3 | KV1-39 | 8 | 4.9 | 0.07 | 0.10 | 0.06 | 0.08 |
| KXD71 | 1 | 4-59 | 16 | 5.8 | LV2-11 | 10 | 5.1 | 0.07 | 0.06 | 0.06 | 0.08 |
| KXD72 | 1 | 3-53 | 9 | 3.4 | KV3-15 | 9 | 4.2 | 0.06 | 0.12 | 0.06 | 0.10 |

Figure S1

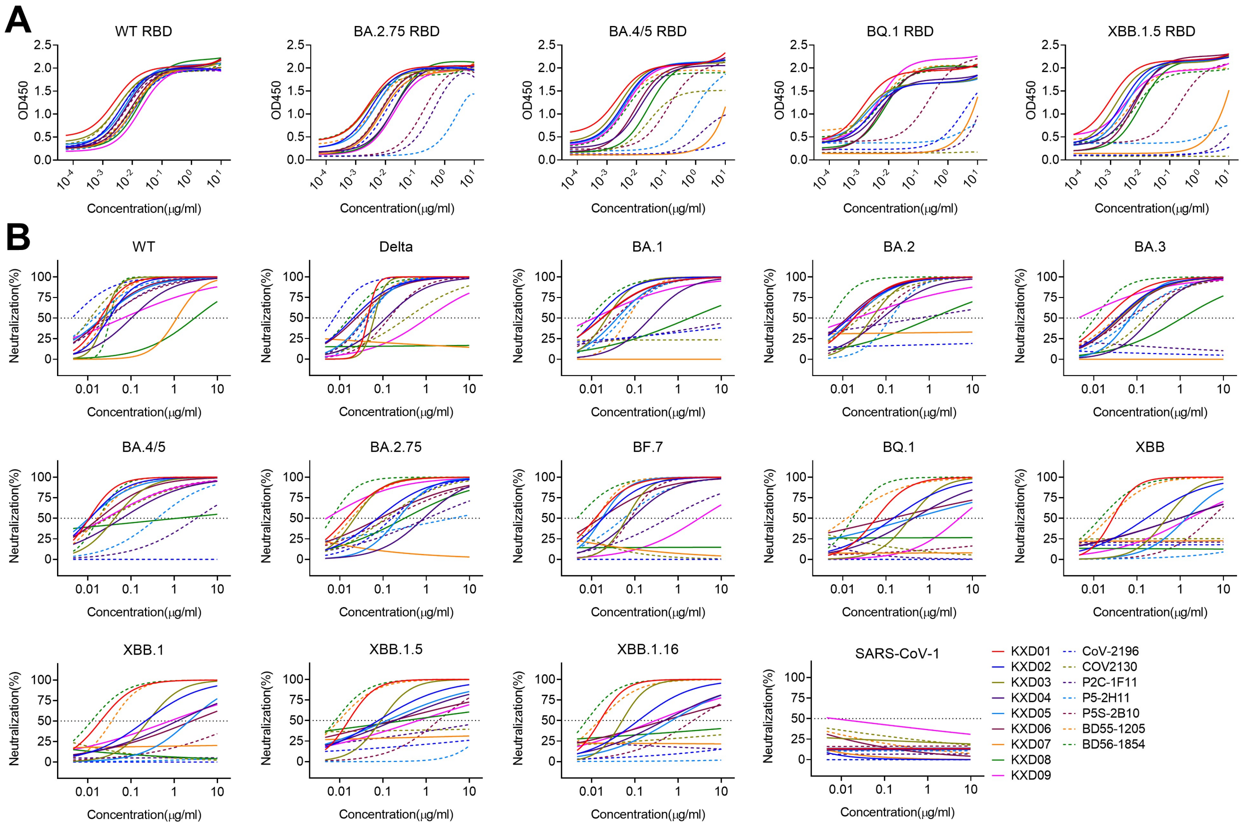
A
WT RBD
BA.2.75 RBD
BA.4/5 RBD
BQ.1 RBD
XBB.1.5 RBD
OD450
Concentration(μg/ml)
B
WT
Delta
BA.1
BA.2
BA.3
BA.4/5
BA.2.75
BF.7
BQ.1
XBB
XBB.1
XBB.1.5
XBB.1.16
SARS-CoV-1
Neutralization(%)
Concentration(μg/ml)
KXD01
KXD02
KXD03
KXD04
KXD05
KXD06
KXD07
KXD08
KXD09
CoV-2196
COV2130
P2C-1F11
P5-2H11
P5S-2B10
BD55-1205
BD56-1854

Figure S2

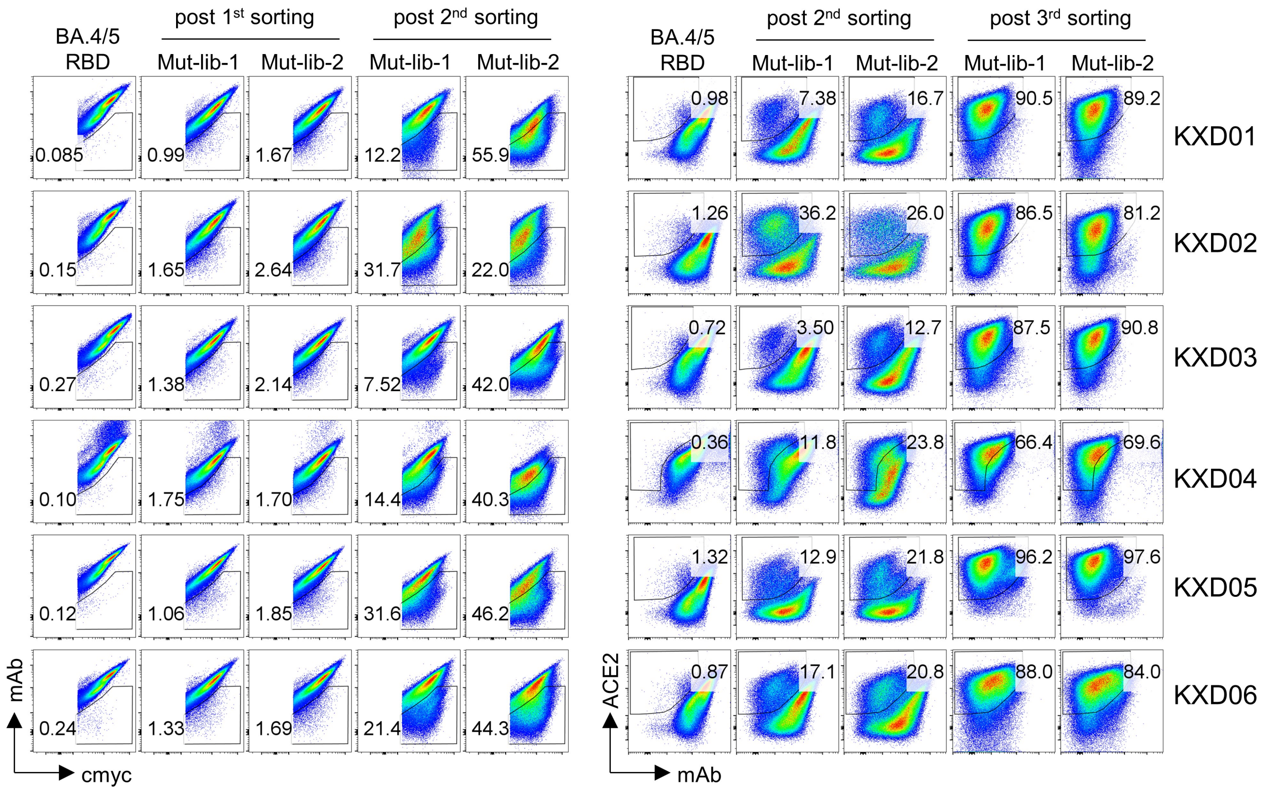
BA.4/5
RBD
post 1st sorting
post 2nd sorting
Mut-lib-1
Mut-lib-2
Mut-lib-1
Mut-lib-2
0.085
0.99
1.67
12.2
55.9
0.15
1.65
2.64
31.7
22.0
0.27
1.38
2.14
7.52
42.0
0.10
1.75
1.70
14.4
40.3
0.12
1.06
1.85
31.6
46.2
0.24
1.33
1.69
21.4
44.3
mAb
cmyc
BA.4/5
RBD
post 2nd sorting
post 3rd sorting
Mut-lib-1
Mut-lib-2
Mut-lib-1
Mut-lib-2
0.98
7.38
16.7
90.5
89.2
KXD01
1.26
36.2
26.0
86.5
81.2
KXD02
0.72
3.50
12.7
87.5
90.8
KXD03
0.36
11.8
23.8
66.4
69.6
KXD04
1.32
12.9
21.8
96.2
97.6
KXD05
0.87
17.1
20.8
88.0
84.0
KXD06
ACE2
mAb

### Figure S3
